# Beyond Normalization: Incorporating Scale Uncertainty in Microbiome and Gene Expression Analysis

**DOI:** 10.1101/2024.04.01.587602

**Authors:** Michelle Pistner Nixon, Gregory B. Gloor, Justin D. Silverman

## Abstract

Though statistical normalizations are often used in differential abundance or differential expression analysis to address sample-to-sample variation in sequencing depth, we offer a better alternative. These normalizations often make strong, implicit assumptions about the scale of biological systems (e.g., microbial load). Thus, analyses are susceptible to even slight errors in these assumptions, leading to elevated rates of false positives and false negatives. We introduce scale models as a generalization of normalizations so researchers can model potential errors in assumptions about scale. By incorporating scale models into the popular ALDEx2 software, we enhance the reproducibility of analyses while often drastically decreasing false positive and false negative rates. We design scale models that are guaranteed to reduce false positives compared to equivalent normalizations. At least in the context of ALDEx2, we recommend using scale models over normalizations in all practical situations.

## Introduction

Sequence count data (e.g., 16S rRNA-seq or RNA-seq data) are ubiquitous in modern biological research. Statistical methods used to analyze these data often fail to control rates of false positives (Hawinkel et al., 2019; Nixon et al., 2023; Roche and Mukherjee, 2022). This phenomenon builds on the broader reproducibility issues in the biomedical sciences (Ioannidis, 2005). Non-biological differences in sequencing depth between samples can substantially contribute to the occurrence of false positives (Gloor et al., 2017; Vandeputte et al., 2017; McGovern et al., 2023; Nixon et al., 2023; Roche and Mukherjee, 2022; Props et al., 2017). In brief, sample-to-sample variation in sequencing depth is often driven by the measurement process, rather than by meaningful variation in the scale (size) of the underlying biological system (Props et al. (2017); Gloor et al. (2017)). To address this problem, many tools incorporate some form of statistical normalization. These normalizations are designed to remove technical variation in sequencing depth so analyses can include between-sample comparisons (Srinivasan et al. (2020)). However, in solving one issue, they can cause another: the choice of normalization can drive inferential results (Nixon et al., 2023; McGovern et al., 2023; Hawinkel et al., 2019; Weiss et al., 2017). Unfortunately, researchers can rarely validate the choice of normalization with real data analyses. Recently, we showed that common normalizations imply modeling assumptions about the unmeasured scale of biological systems (Nixon et al., 2023). We found false positive rates as high as 80% with only slight errors in these implicit assumptions.

To study this scale issue, we formulated the problem using partially identified statistical models and found simple, intuitive conclusions (Nixon et al., 2023). When a research question requires knowledge of the system scale but the observed data lacks that information, researchers need to make some modeling assumptions about the system scale. For instance, in differential abundance analysis using 16S rRNA gene-sequencing, scale assumptions are often made implicitly through the chosen normalization. However, these assumptions should be an explicit part of the model-building process to enhance the transparency and reproducibility of research. Moreover, statistical methods must incorporate potential errors from scale assumptions to make resulting analyses rigorous. To facilitate such analyses, we introduced a specialized and computationally efficient family of Bayesian partially identified models, called *Scale Simulation Random Variables* (SSRVs). Rather than using a single normalization, SSRVs use a *scale model* to represent uncertainty in the scale of the underlying system. Expert knowledge alone can specify scale models, in which case they generalize standard normalizations, or they can be models of external scale measurements (e.g., qPCR). Moreover, SSRVs are more flexible than prior methods which use sparsity assumptions (e.g. Grantham et al. (2020)) because scale models can be designed based on sparsity assumptions, but such assumptions are not required. Through analysis of both real and simulated data, we demonstrated that accounting for scale uncertainty as part of modeling can dramatically reduce both Type-I (false positive) and Type-II (false negative) error rates (Nixon et al., 2023; McGovern et al., 2023).

This article presents an intuitive introduction to scale model-based analysis and an update to the popular ALDEx2 library. First, we illustrate the problem with existing normalizations using an explicit example. Next, we review the ALDEx2 library and the origin of its implicit scale assumptions. We describe our updates to the ALDEx2 library which allows users to replace normalizations with scale models, making it the first general-purpose suite of tools for scale model analyses. We highlight a class of scale models that generalize the prior normalizations of ALDEx2 and are guaranteed to reduce false-positive rates. Through four case studies, we demonstrate that scale model analysis can drastically decrease false positive rates. In multiple studies, scale model-based analyses control false positive or false discovery rates at nominal levels (e.g., 0.05%) when normalization-based methods such as DESeq2 (Love et al., 2014), edgeR (Robinson et al., 2010), baySeq (Hardcastle and Kelly, 2010), and limma (Ritchie et al., 2015) all display rates above 50%. Moreover, by designing scale models based on flow-cytometry data or biological knowledge, we show that scale model analysis can also reduce false negative rates. Due to their remarkable performance improvements and our prior theoretical work, we recommend using scale models in ALDEx2 rather than normalizations in all practical situations.

## An Illustration of the Problem with Normalizations

Figure 1 illustrates two microbial communities: one control condition and one treated with a drug of interest. Samples are obtained and sequenced from both communities to estimate the effect of the drug. After sequencing, 2,000 reads mapped to a particular taxon in Community A (square taxon), whereas 5,000 reads mapped to that same taxon in Community B. From this information, it appears the drug is associated with an *increase* in the taxon’s abundance. However, this is not necessarily the case: sequencing depth can alter our conclusion. Here, Community B was sequenced deeper (10,000 total reads) than Community A (3,000 total reads). Typically, one uses normalization to remove this confounding technical variation, giving the measurements a common scale.

**Figure 1:**
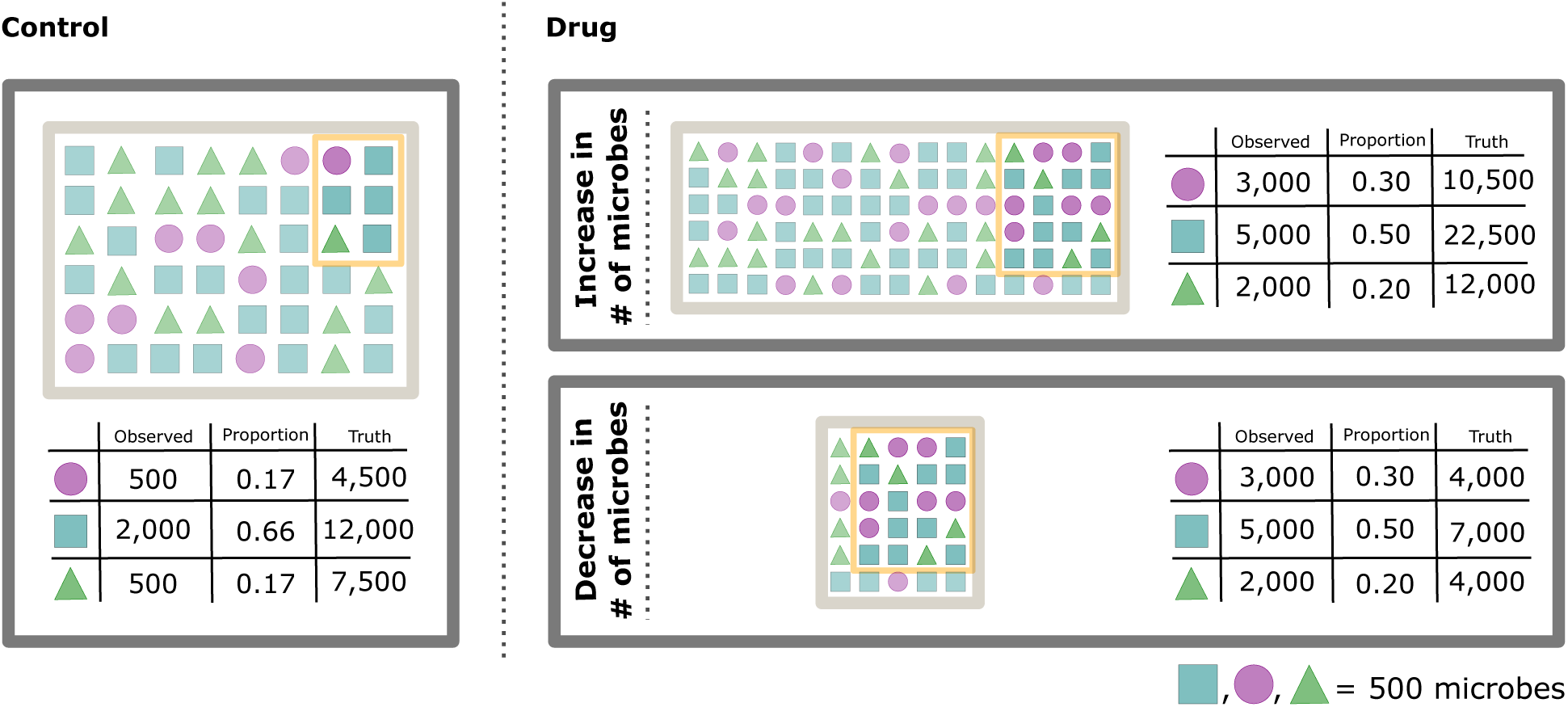
Scale can confound sequence count data analyses. In this study, we measure counts of three different types of microbes (square, circle, triangle) from samples of a control community and a drug-treated community. The yellow boxes represent our samples (“Observed” columns), while the larger gray boxes represent the entire community (“Truth” columns). In the drug condition, the same observed sample could have come from a large community (top box) or a small community (bottom box). Total Sum Scaling (TSS) normalization estimates the proportional abundance of each type of microbe in the underlying system (“Proportion” columns). However, these proportions are insufficient to determine if a particular taxon increases or decreases in abundance in response to the drug.

With Total Sum Scaling (TSS) normalization, we transform the observed data to proportional amounts by dividing the observed counts by the sequencing depth as illustrated in the “Proportion” columns in Figure 1. The normalized measurements have a common scale: they each sum to one. Based on these proportions, it appears the drug is associated with a *decrease* in the abundance of the square taxon.

At first glance, it seems that normalization has corrected our previous erroneous conclusion, but, unfortunately, TSS normalization is insufficient to determine if the square taxon increases or decreases in response to the drug. In drawing conclusions about the underlying communities from those TSS normalized measurements, we implicitly assumed a constant scale of the underlying communities. That is, we implicitly assumed that the drug and control communities have the exact same microbial load. But, if the scale of the underlying system was larger in Community B than Community A (top right box of Figure 1), then this taxon would have *increased* in the drug case. Figure 1 shows that the taxon could, in truth, be either increasing or decreasing depending on whether the microbial load in the drug condition is higher or lower than in the control. This disparity arises because TSS normalization does not take into account the scale of the underlying system.

This illustration is an example of differential abundance or expression (DA/DE) analysis, which investigates if any of the *D* entities are present in different amounts (different abundances or different levels of expression) in two biological conditions (here, control and drug). This goal is often formalized as a problem of estimating log-fold changes (LFC). The LFC of each entity *d* is defined as the difference in the average log-transformed amounts of the entity between the two conditions: 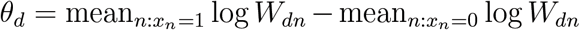 where *x*_*n*_ denotes the condition for sample *n* and *W*_*dn*_ denotes the amount of entity *d* in biological system *n*. For example, in Figure 1, the LFC of the square taxon is log_2_ 22, 500 *−* log_2_ 12, 000 = 0.90 in the increased load case (top box) and log_2_ 7, 000 *−* log_2_ 12, 000 = *−*0.78 in the decreased load case (bottom box).

As in our illustration, common tools that use normalization do not estimate LFCs with the true amounts (*W*_*dn*_), but instead use normalized amounts. For example, using TSS normalized amounts, we estimate the LFC of the square taxon as log_2_ 0.5 *−* log_2_ 0.66 = *−*0.40 regardless of whether the drug condition has increased or decreased load. In Supplementary Section S.1, we show that the LFC estimate from TSS normalized data is only correct if the microbial load is exactly equal in the two conditions: 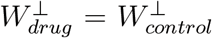 where 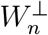 denotes the total microbial load in system *n*. If this condition is unmet, then the estimated LFC will be biased, and resulting hypothesis tests may display elevated rates of false positives, false negatives, or both.

This example illustrates an uncomfortable fact: controlling for technical variation in the scale of data differs from recovering biological variation in the scale of systems. To address this deficiency, some authors supplement sequence count data with external measurements of the system scale, e.g., qPCR, flow cytometry, or DNA spike-ins (Vandeputte et al., 2017). While these methods can help, they are not a universal solution for at least three reasons. First, either due to cost or effort, researchers infrequently collect external measurements; hence, public datasets often lack those data. Second, even if those measurements are collected, they can be noisy and often require specialized statistical methods (Galazzo et al., 2020). Finally, these methods may not measure the relevant scientific scale. For example, in studying human microbiota, DNA spike-ins added after DNA extraction may recover variation in DNA concentration within DNA libraries (Stämmler et al., 2016). However, variation within the DNA libraries may still differ from variation in microbial load within the human gut.

## Results

### From Normalizations to Scale Models in ALDEx2

ALDEx2 is a popular tool for DA/DE analysis (Fernandes et al., 2014). While ALDEx2 can perform a wide range of log-linear modeling tasks, we focus on it as a tool for DA/DE analysis and leave a discussion of its more general linear modeling capabilities to *Supplemental Materials*. Here, we briefly describe how it works; a more formal description can be found in *Methods*.

First, ALDEx2 uses a Bayesian model to simulate proportional amounts, taking into account the randomness of the sequencing process. Second, ALDEx2 uses a Centered LogRatio (CLR) transform to normalize the estimated proportions. Third, ALDEx2 estimates LFCs using the CLR normalized amounts. Finally, for each entity, a summary p-value is calculated for a test of the null hypothesis that the LFC of the entity equals zero (no DA/DE). While there were several technical details we needed to address (see Supplementary Sections S.2-S.4 for details), our principal modification of ALDEx2 is a change to the second step.

Like TSS normalization, the use of the CLR normalization makes an implicit assumption about the system scale. Let 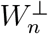 denote the scale of the *n*-th system and 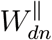 the proportional amount of the *d*-th entity in that system. In *Methods*, we show the CLR normalization corresponds to an assumption that 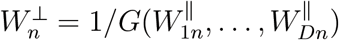 where *G* denotes the geometric mean function. That is, ALDEx2 assumes that the system scale can be imputed without error from the proportional amounts of each entity, an assumption contradicted in our illustrative example.

Normalizations like the CLR and TSS have two critical limitations. First, their assumptions about the system scale are implicit and often unrecognized by researchers. Second, these assumptions are strict: resulting LFC estimates and statistical inferences are only valid if the assumptions hold exactly (Nixon et al., 2023). An intuitive solution to both problems is to make these assumptions an explicit part of the model-building process, then deal with potential errors that arise from those assumptions. Consider a model which generalizes the CLR normalization assumption^1^:

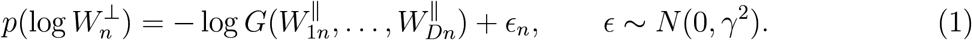

When *γ*^2^ *→* 0, then *ϵ*_*n*_ *→* 0 and this model is equivalent to the assumption underlying the CLR normalization (on a log-scale 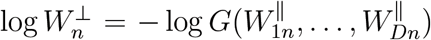). However, when *γ*^2^ *>* 0, the model allows for potential error in that assumption. This is an example of a *scale model*. More generally, a scale model is any probability model for the scale of the system: 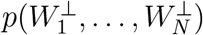. By accounting for potential error in normalization assumptions, scale models can drastically reduce false positives (Nixon et al., 2023). Moreover, scale models are flexible and allow analysts to specify more biologically realistic assumptions than off-the-shelf normalizations, thereby reducing false negative rates (Nixon et al., 2023). Finally, scale models can be simple; in *Methods*, we discuss various features of DA/DE analysis that reduces the burden in scale model specification. We demonstrate these and other features of scale models in later sections.

While there were several technical details that we overcame to implement scale models within the ALDEx2 software library (see Supplementary Sections S.2-S.4 for details), our overarching approach is intuitive. In the following, a 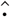 denotes an estimate. In prior versions of ALDEx2, each estimate of the systems’ proportions 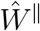 was normalized by a function (e.g., the CLR transform) to an estimate of the absolute amounts: 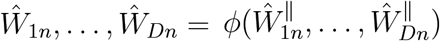. We replace this step with a sample from a scale model 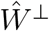. For each simulated estimate of proportional abundances 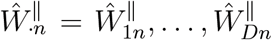, we produce a corresponding sample from the scale model 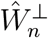. Multiplying these together produces an estimate of the absolute amounts:

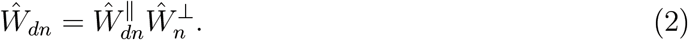

That is, absolute amounts 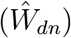 are equal to proportional amounts 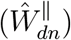 times scale 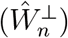. The resulting estimates 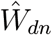 can be used in subsequent steps of the ALDEx2 model just as before. In practice, this algorithm has no perceptible increase in runtime or memory demands compared to the original ALDEx2 software.

### Scale Models Can Dramatically Decrease False Discovery Rates

To illustrate the advantages of scale models, we reproduced the simulation study of Nixon et al. (2023) (*Methods*). We simulated the true abundance of 20 taxa in two conditions (preand post-treatment with a narrow-spectrum antibiotic). After treatment, 4 of the 20 taxa decrease in abundance. We simulated the lack of scale information in the observed data by resampling the true abundances to an arbitrary sequencing depth. We benchmarked the resampled data using standard tools for DA/DE analysis including the original ALDEx2 model (with CLR normalization), DESeq2 (Love et al., 2014), edgeR (Robinson et al., 2010), baySeq (Hardcastle and Kelly, 2010), and limma (Ritchie et al., 2015) (Figure 2). All of these methods were unreliable and, at the largest sample sizes, demonstrated over three-times more false positives than true positives. More concerning, the false positive rate for all these methods increased to over 75% with larger sample sizes (Figure 2). This result contradicts standard statistical wisdom: inferential performance is supposed to improve with more data.

**Figure 2:**
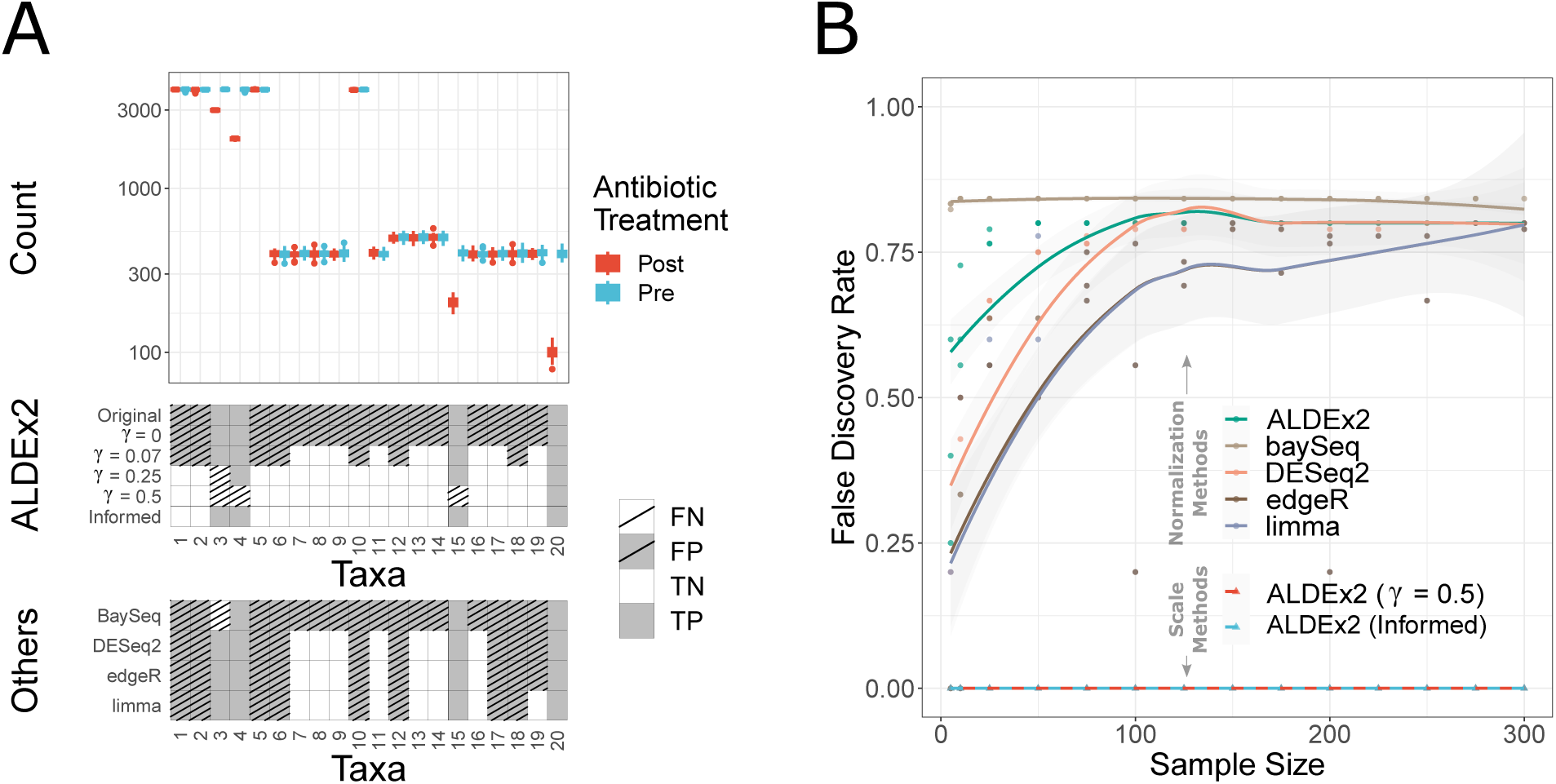
Scale models can drastically decrease false positive rates. The true abundances of 20 microbial taxa were simulated before and after treatment with a mild, narrow spectrum antibiotic (*Methods*). **A**. The top panel (“Count”) shows simulated true counts (N=50 per condition) for each of the 20 taxa: only taxa 3, 4, 15, and 20 change between conditions. The bottom two panels (“ALDEx2” and “Others”) shows the true positives (TP), false positives (FP), true negatives (TN) and false negatives (FN) for ALDEx2 and many common methods applied to the resampled data. We compared the original normalizationbased ALDEx2 model (“Original”) to the default scale model for several values of *γ* and an Informed model mimicking a slight decrease in microbial load after antibiotic administration. **B**. The same simulation in Panel A was repeated in triplicate over data sizes ranging from 5 to 300 samples per condition. Only the scale-based ALDEX2 models [ALDEx2 (*γ* = 0.5) and ALDEx2 (Informed)] control false discovery rates asymptotically.

**Figure 3:**
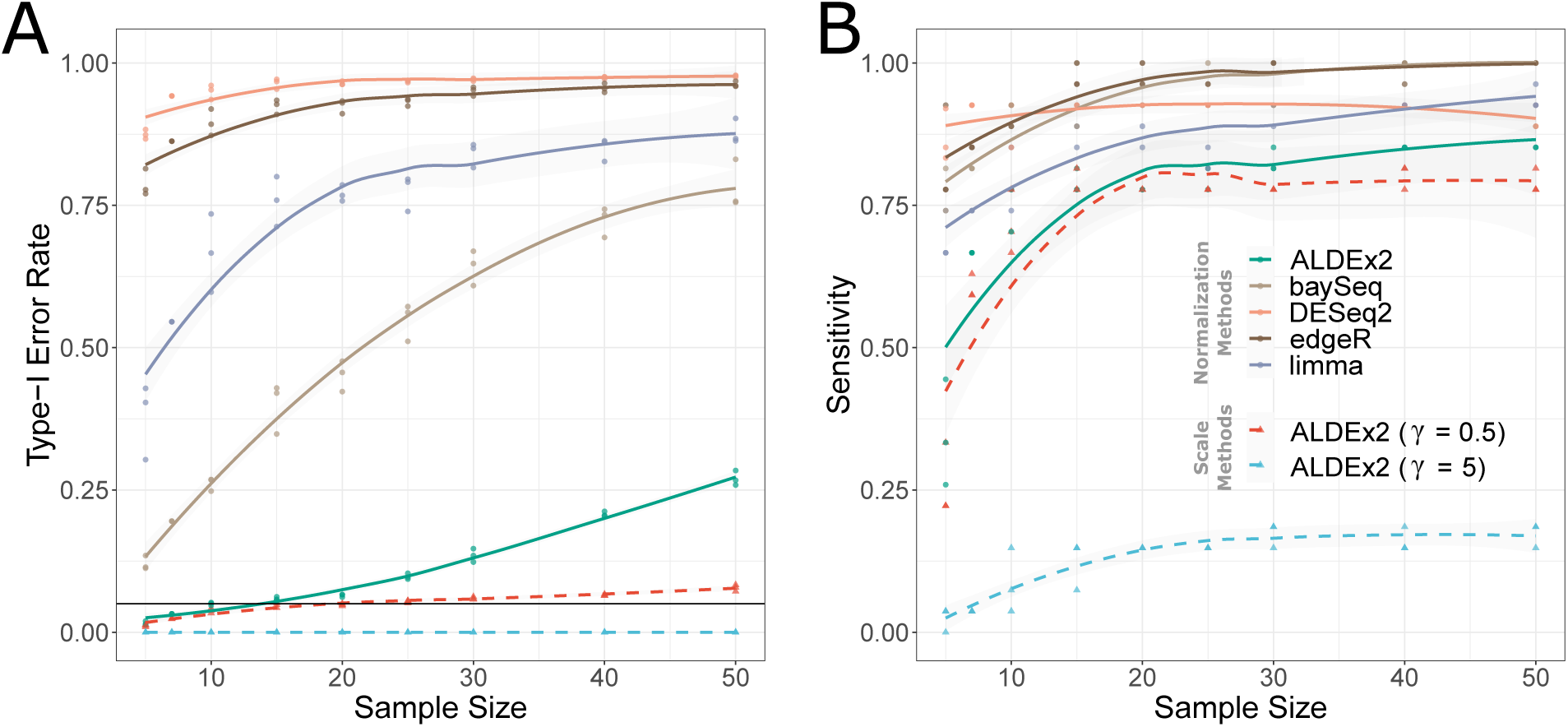
Incorporating Scale Uncertainty Improves Performance over Normalizations in a Selective Growth Experiment (SELEX). We reanalyzed the SELEX study at different sample-sizes with data resampling (see main text and *Methods*). For each resampled dataset, we applied ALDEx2 with the default scale model (with *γ* = 0.5, 5 ). We also applied five other normalization-based methods commonly used differential expression analysis. **A**. Type-I error rates for each tested method. We applied a mean-based smoother for better visualization. For each method, statistical significance was determined at a threshold of *α* = 0.05 based on multiplicity-adjusted p-values. Therefore each method should control type-I error at or near a level of 0.05 (black horizontal line). Only methods that account for scale uncertainty achieve this for all sample-sizes. **B**. The sensitivity for each method with mean-based smoother. While many methods have high sensitivity, they do so with a high rate of false positives. Yet, even at extreme levels of scale uncertainty *γ* = 5, ALDEx2 identifies true positives while controlling the number of false positives.

This bizarre phenomenon, where type-I errors increase with more data, is a hallmark of an *unacknowledged bias* (Nixon et al., 2023; Gustafson, 2015). The LFC estimates produced by these methods are biased due to errors in their implicit assumptions about scale. These methods fail to consider such errors; as the sample size increases, these methods become increasingly confident in their incorrect (biased) estimate. Scale models can mitigate this problem by allowing researchers to consider errors in these assumptions or even make more biologically plausible assumptions.

*Our new default scale model* for ALDEx2 demonstrates how considering errors in modeling assumptions can improve inferences. To stay consistent with prior versions which used the CLR normalization, the default scale model generalizes the CLR transform, as did Equation (1). The default scale model has better asymptotic performance than Equation (1) (see *Methods*). Like Equation (1), the default scale model includes one user-defined parameter *γ* which controls the amount of uncertainty in the CLR assumption. When *γ* = 0, we recover the original ALDEx2 model; when *γ >* 0, we account for error in the CLR normalization assumption. In fact, for any value of *γ >* 0, the new ALDEx2 model will provide better type-I error control than the original ALDEx2 model. As a general guideline, we recommend *γ* = 0.5 as a reasonable default value for most cases (see *Methods* for explanation based on the interpretation of *γ*). Figure 2 demonstrates that by incorporating even tiny amounts of scale uncertainty (*γ >* 0.07), the false positive rate of ALDEx2 drops precipitously while still revealing true positives.

In Supplementary Section S.5, a sensitivity analysis shows how the choice of *γ* influences false positive and false negative rates. For any biologically reasonable choice of *γ* (see discussion in that supplementary section), the new ALDEx2 model controls the false discovery rate while simultaneously revealing true positives. That section also discusses how sensitivity analyses can facilitate novel and transparent forms of reporting and can sometimes even eliminate the need to choose a single value of *γ*.

We next designed an *informed* scale model based on our knowledge that the antibiotic is narrow spectrum; we expect only a small decrease in the microbial load (see *Methods*). Compared to the default scale model, this informed model reduces the bias of LFC estimates by reflecting more biologically reasonable beliefs. While the default scale model reduced false positives compared to the original ALDEx2 model, the informed scale model also reduces false negatives (Figure 2).

Figure 2B shows that of all the methods tested, only those that used scale models mitigated unacknowledged bias and control false discovery rates at a nominal 0.05% as sample sizes increased. All other methods displayed false discovery rates above 75% when given enough data.

### Scale Uncertainty Enhances the Reanalysis of a Selective Growth Experiment

We reanalyzed the Selective Growth Experiment (SELEX) study originally published in McMurrough et al. (2014) and later highlighted in the original publication of ALDEx2 (Fernandes et al., 2014). Researchers wanted to identify which of 1,600 gene variants conferred a growth phenotype upon cell lines exposed to a bacteriostatic toxin. They designed the study so that cell lines with variants capable of removing a toxin increased in abundance when exposed to the toxin while all other cell lines remained unchanged. They took samples from two experimental conditions, one with (selective) and one without the toxin (non-selective). This dataset is useful for two reasons. First, some of the variants have been verified *in vitro*, giving an objective measure of truth. Second, the directionality of abundance changes between conditions was asymmetric and fixed: any changes in cell line abundance were increases in the selective (rather than non-selective) growth condition. Thus, we can use biological knowledge to design an appropriate scale model.

For this data, the CLR normalization makes the implicit assumption that the absolute scale is approximately 265 times higher in the selective (bacteriostatic toxin) versus nonselective (control) condition (see *Methods*). Our biological knowledge supports this direction of change, but the magnitude of the change is uncertain. We use the default scale model to express uncertainty in the CLR assumption.

We used repeated data sub-sampling to investigate how the type-I error (false positive error) rates and sensitivity of different methods varied as a function of sample size. Based on validation experiments (*Methods*), we knew that only a small fraction (27/1,600) of genes confer a growth-promoting phenotype. However, existing methods returned many more genes as significant (e.g., around 1,500 genes are returned as significant by DESeq2 at a sample size of 10). The only methods capable of controlling type-I errors are the scale-based ALDEx2 models. At the generally recommended value of *γ* = 0.5, ALDEx2 provides loose Type-I error control: type-I error only increases above the stated 0.05 level for the largest sample sizes yet still remains near the stated level. We also tested ALDEx2 with an unreasonably large value of *γ* = 5 to illustrate performance even with over-estimated uncertainty. At *γ* = 5, ALDEx2 displays zero false positives for any sample size and still identifies five true positives. Still, at *γ* = 5, sensitivity is lower than that of *γ* = 0.5: in the latter, case the sensitivity of ALDEx2 is comparable to other methods with only a fraction of the false positives of other methods.

### Informative Scale Models Can Reduce False Negatives

The prior two sections showed that false positives can be drastically reduced by integrating potential error in assumptions about scale. Here, we show how false negatives can decrease when scale models better reflect biology.

We reanalyzed a recent study by Vandeputte et al., who proposed supplementing 16S rRNA microbiome data with flow-cytometry based measurements of fecal microbial concentration (Vandeputte et al., 2017). This study compared fecal microbiota between 29 patients with Crohn’s Disease (CD) and 66 healthy controls. The original authors analyzed these data using a method they called Quantitative Microbiome Profiling (QMP): first, they rarefied the sequence count data to an even sampling depth, and then they multiplied the rarefied counts by the measured flow-cytometry measurements. In the present context, this can be thought of as using Equation (2) without considering measurement noise in the composition or scale. In contrast, we can account for measurement noise in the sequence count data and in the flow-cytometry by using ALDEx2 with a flow-cytometry-informed scaled model:

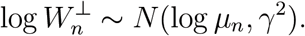

In this model, *μ*_*n*_ denotes the flow-cytometry measurement for total cellular concentration in the *n*-th fecal sample, and *γ* is related to the error in that measurement technique (see *Methods*). Treating the results of this model as our gold-standard, we benchmarked a variety of other DA/DE tools, including QMP and ALDEx2 with different scale models.

Echoing the results of Vandeputte et al. (2017), normalization-based methods (including the original ALDEx2) demonstrate elevated rates of both false-positives and false negatives (Figure 4A). Remarkably, QMP missed three bacterial genera that are differentially abundant between groups: *Parabacteroides, Sutterella*, and *Anaerobutyricum*. All three of these genera have been previously associated with Crohn’s disease (Wang et al. (2018); Cui et al. (2022); Suskind et al. (2020)). Moreover, Figure 4B shows that our conclusions about these three taxa are largely insensitive to flow-cytometry measurement error. We suspect all three of these false negatives arise from rarification, which can lead to decreased statistical power by throwing away data (McMurdie and Holmes, 2014).

**Figure 4:**
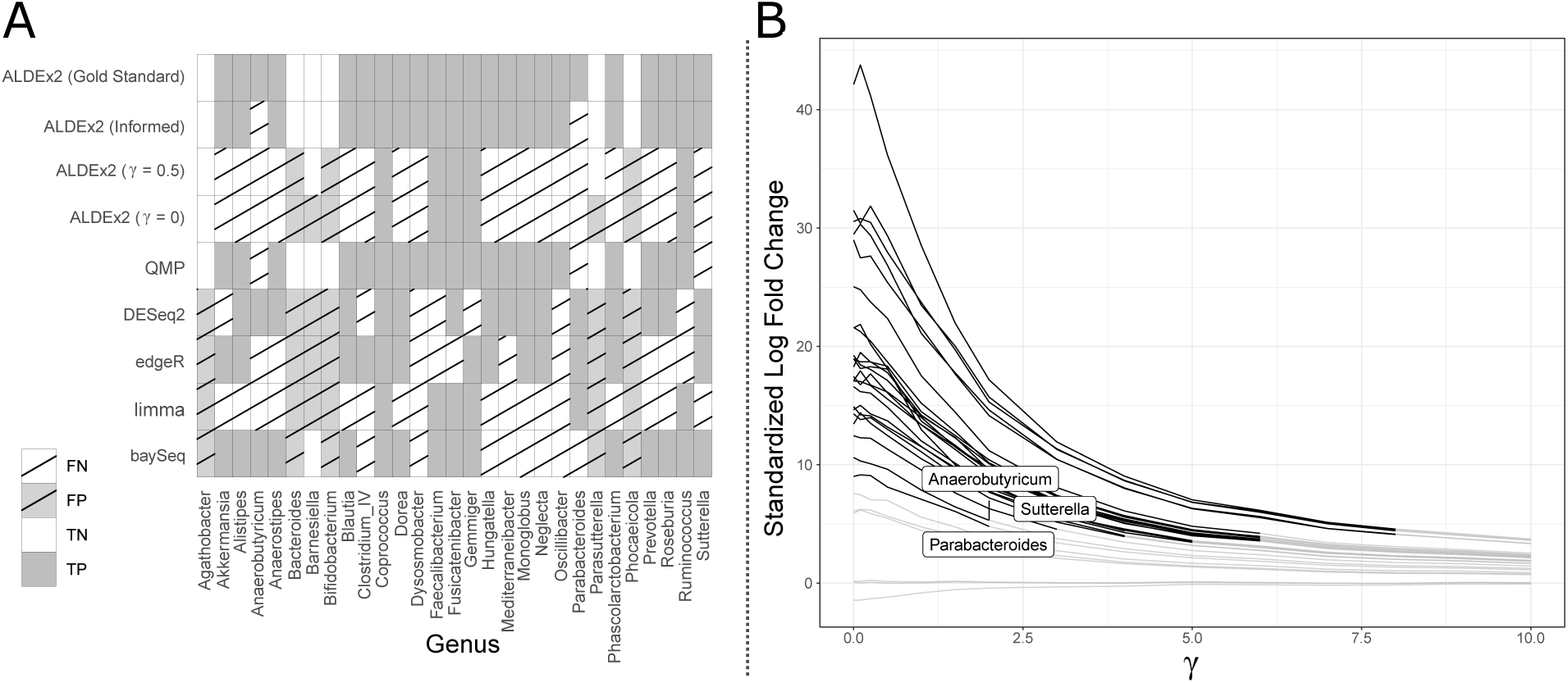
Scale Models Built from Outside Measurements or Biological Reasoning can Reduce Both Type-I and Type-II Error Rates. **A**. The pattern of false positives (FP), false negatives (FN), true positives (TP), and true negatives (TN) for each differential abundance tool applied to the Vandeputte et al. (2017) study data. Only taxa identified as differentially abundant by at least one method are shown. True/False Positives/Negatives were defined based on ALDEx2 with a *Gold Standard* scale model which integrated flowcytometry measurements of total microbial concentration in fecal samples (see *Methods*). The QMP model also had access to these measurements but did not account for compositional uncertainty or measurement error in the flow-cytometry measurements. The *Informed* scale model was based on visual inspection of an independent study which used real-time PCR to quantify microbial load in healthy versus Crohn’s patients (see *Methods*). *γ* = 0 and *γ* = 0.5 refer to the default scale model in ALDEx2. **B**. Sensitivity analysis showing how the standardized log fold change (the average LFC across Monte Carlo samples divided by the standard deviation across Monte Carlo samples) varies with different levels of measurement error in the flow-cytometry measurements. Each line corresponds to a single taxon and is grey if *p >* 0.05 and black if *p ≤* 0.05. For reference, the Gold Standard scale model in panel A uses *γ* = 1 based on data available in Vandeputte et al. (2017) (*Methods*). We label the three taxa identified by ALDEx2 with the Gold Standard scale model but not QMP.

Both QMP and our flow-cytometry informed ALDEx2 model decrease false positive and false negative rates by using supplemental measurements of microbial concentration. As these measurements are often unavailable, we evaluated whether we could design an equally effective *Informed* scale model based only on visual inspection of figures present in an independent study of Crohn’s disease (Sarrabayrouse et al., 2021). Like Vandeputte et al. (2017), Sarrabayrouse et al. (2021) estimated total microbial load in patients with CD compared to healthy controls. Unlike Vandeputte et al. (2017), which studied an Belgian cohort with flow-cytometry for microbial load measurements, Sarrabayrouse et al. (2021) studied a Spanish cohort with quantitative PCR measurements and validated their findings on a Belgian cohort. Biases due to copy-number variation or DNA extraction could make these measurement techniques incomparable. Despite these differences, our *Informed* scale model, built by visual inspection of Figure 2 of Sarrabayrouse et al. (2021) (see *Methods*) provided nearly identical results to QMP (Figure 4A; type-I and type-II error rates of 0% and 9% respectively). This result highlights that even weak expert knowledge about scale, when expressed as a scale model, can enable dramatic decreases in both false positive and false negative rates compared to normalization-based methods.

We evaluated the CLR normalization and the default scale model in more detail. In short, the CLR normalization does poorly in this case study: the CLR here equates to an assumption that microbial load in CD patients is substantially increased compared to healthy controls when the results of Vandeputte et al. (2017) and Sarrabayrouse et al. (2021) suggest a slight decrease. These results emphasize the importance of interrogating assumptions about scale as part of sequence count data analyses. If a researcher did want a CLR assumption to analyze these data, they would improve inference by considering scale uncertainty: the default scale model with the default value of *γ* = 0.5 achieves the same sensitivity as original ALDEx2 model with the CLR normalization yet has fewer false positives.

### Scale Uncertainty Can Lead to More Biologically Plausible Results in RNA-Seq Studies

As a final case study, we reanalyzed a RNA-seq study which has been use to inform sample size selection for gene expression studies (Gierliński et al., 2015; Schurch et al., 2016). This study contained 48 biological replicates from each of two *Saccharomyces cerevisia* strains: a wild-type (WT) and SNF2-knockout (SNF2) strain. As reported by Schurch et al. (2016), existing tools often report over 70% of genes are differentially expressed in response to this knock-out. We hypothesized that this percentage included a large number of false positives arising from errors in scale assumptions. In Supplementary Section S.6, we describe this analysis and show that many of these differentially expressed genes are no longer significant when one accounts for even small amounts of scale uncertainty. For example, with ALDEx2’s default scale model, in moving from *γ* = 0 to *γ* = 0.25, the proportion of genes identified as differentially expressed drops from 68% to just 12%. While we lack a ground truth measure of what genes are differentially abundant, these results suggest that normalization-based methods may have substantial inflation of type-I errors due to a lack of uncertainty in scale assumptions.

## Discussion

While implicit modeling assumptions may bias results in the analysis of sequence count data, the choice of normalization can dominate model estimates and obscure biological conclusions (Nixon et al., 2023; Clausen and Willis, 2022; Weiss et al., 2017; Props et al., 2017; Vandeputte et al., 2017). Here, we introduced scale models as a generalization of normalizations, so researchers can account for potential errors in their implicit modeling assumptions about scale. We introduced the updated ALDEx2 software package, which provides the first general-purpose suite of tools for scale model analysis. Through case studies, we showed that accounting for potential errors in scale assumptions can drastically reduce false positive rates. Beyond generalizing normalizations, we showed that scale models can be built from prior knowledge or external scale measurements. By better reflecting biology, such scale models can also reduce false negatives.

Previous research has suggested generalizing beyond a single normalization in analyzing sequence count data. For example, Song et al. (2023) introduced a method of combining p-values obtained under different normalizations into an overall p-value. However, such work has fundamental limitations– the assumptions underlying normalizations are often implicit, obscuring which normalizations cover the biologically plausible range of assumptions. There may also be cases where none of the available off-the-shelf normalizations adequately cover actual biology. In contrast, the assumptions underlying scale models are explicit, and standard probability tools can customize scale models to any given study.

Outside of analyzing sequence count data, our work connects to the topic of rigor and reproducibility in statistics and machine learning. To address reproducibility problems in science (e.g., Ioannidis (2005)), some authors have suggested increased attention on *stability* : conclusions drawn from data should withstand perturbations of the observed data and the chosen model. For example, in computer vision, researchers often include perturbed data (e.g., rotated images or Gaussian noise) to reduce over-fitting and help models generalize beyond the training set. More recently, some authors have suggested that statistical inference should integrate similar ideas (Yu, 2020). Our work improves the stability of ALDEx2: the scale model accounts for perturbations to the chosen normalization. Under the new ALDEx2 model, reported p-values and confidence intervals now include a stability guarantee: conclusions drawn based on these quantities are robust to the perturbations encoded in the scale model.

This work demonstrated how to improve the rigor of existing tools by accounting for scale uncertainty. We minimized our changes to ALDEx2 to highlight that scale uncertainty, rather than other changes, drives observed performance improvements. Many avenues exist for future refinement, including developing new scale models. While we included the default scale model which is reasonable for many cases, it is not a universal solution. Although the default scale model can reduce false positives compared to the CLR normalization, scale models that better reflect biology may further decrease in false positives *and* false negatives (Nixon et al., 2023).

Fortunately, scale models can be incorporated into some other modeling software. For example, in Nixon et al. (2023), we created SSRVs by combining regression models for the system composition built from the *fido* software package with scale models (Silverman et al., 2022). If a tool models system composition, we can incorporate scale models into it in the same manner: identify the relevant system compositing parameter, multiply it by samples from a scale model, and create an SSRV. We expect researchers can use this method to extend *Songbird* (Morton et al., 2019), *GPMicrobiome* (Ä ijö et al., 2018), *PhILR* (Silverman et al., 2017), and *propr* (Quinn et al., 2017) *to create scale models for scale reliant inference. Unfortunately, beyond these tools, the picture is less clear. For example, we have yet to identify how to incorporate scale uncertainty into tools like DESeq2* (Love et al., 2014), *edgeR* (Robinson et al., 2010), *or limma* (Ritchie et al., 2015), *which lack parameters that can be identified with the system composition*.

## Methods

### Problem Set-Up and Notation

*We denote a sequence count dataset as a D × N* matrix *Y*, with elements *Y*_*dn*_ denoting the number of sequenced DNA molecules mapping to the *d*-th entity (e.g., taxa or gene) in the *n*-th sample. Following Nixon et al. (2023), we think of the observed data as an imperfect measurement of an underlying biological system *W* called a *scaled system*. We represent the scaled system *W* as a *D×N* matrix whose elements *W*_*dn*_ represent the true amount of entity *d* in the biological system from which the *n*-th sample was taken. The notion of *true amount* depends on both the studied system and the scientific question, e.g., the true amount could represent bacterial cell count, colony-forming units (CFUs), or cellular concentration in a medium in microbiota studies

The term *scaled system* alludes to the fact that *W* can be uniquely described in terms of its scale (i.e., summed amounts, *W*^*⊥*^) and composition (i.e., proportional amounts, *W*^*∥*^) via:

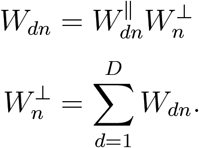

These relations imply *W*^*∥*^ is a *D × N* matrix with columns summing to one 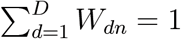, e.g., the columns of W are compositional vectors) while *W*^*⊥*^ is an *N* -vector. When we say that sequence count data (*Y* ) lacks information about the system scale, we are referring to the fact that sample-to-sample variation in sequencing depth (i.e., 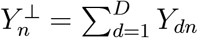) is driven by the measurement process; such variation is typically unrelated to meaningful biological variation in the scale of the system 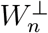 (Props et al., 2017; *Vandeputte et al., 2017)*.

*We use a* 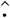 to distinguish between an estimate of a quantity and its corresponding true value (e.g., 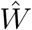 vs *W* ). When working with samples of a quantity obtained via computer simulation, we use super-script ^(*s*)^ to denote the *s*^*th*^ sample. When a quantity depends on only composition or scale, we use a superscript ^*∥*^ and ^*⊥*^, respectively. Finally, when discussing rows or columns of a matrix we use a subscript “*·*”, e.g., *W*_*·n*_ refers to the *n*-th column of the matrix *W* .

### The Normalization-Based ALDEx2 Model

The ALDEx2 model consists of four main steps (Fernandes et al., 2014). First, *S* samples of the system composition are drawn from the posterior of *N* independent multinomial-Dirichlet models. The posterior of the *n*-th model is given by:

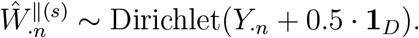

where **1**_*D*_ denotes a *D*-length vector of 1s. Each posterior sample is then normalized using one of several built-in normalizations (the default is the CLR). Each normalization can be expressed as a sample-wise transformation:

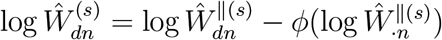

where *ϕ* is defined by the chosen normalization. These normalized samples are then used to estimate log-fold-changes (LFCs) for each entity *d*

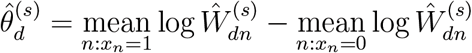

where *x*_*n*_ *∈ {*0, 1*}* is a binary variable denoting the two conditions (e.g., disease versus health). A parametric or non-parametric test then examines the null hypothesis that 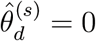. Finally, ALDEx2 summarizes over the *S* samples, reporting the mean p-value and LFC estimate for each entity. See Supplementary Section S.3 for a more formal definition of the ALDEx2 model and details of its linear modeling capabilities.

### Scale Assumption implied by CLR Normalization

ALDEx2’s normalization step introduces an assumption about the system scale. Note that the decomposition 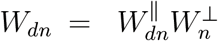 presented in the problem set-up can be equivalently stated as 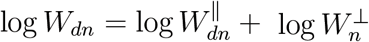. Comparing this to the normalization equation:

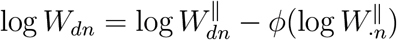

reveals the assumption that

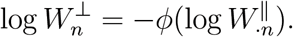

This assumption says that the system scale can be imputed, without error, as some known function of the system composition. The centered log-ratio (CLR) normalization is defined by 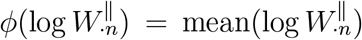, which implicitly assumes that the scale of the system is related the geometric mean of the composition

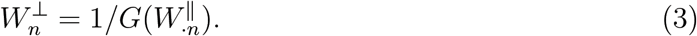

### A New, Scale-Based ALDEx2 Model

A scale model is a probability model for the system scale: 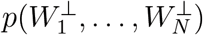. The new version of the ALDEx2 model replaces normalizations with a scale model. As before, it still samples from the system composition *W*^*∥*(*s*)^. But rather than normalizing *W*^*∥*(*s*)^ to estimate the system *W* ^(*s*)^, ALDEx2 now samples from a scale model and multiplies the composition (*W*^*∥*(*s*)^) by that sample (*W*^*⊥*^) to estimate the system:

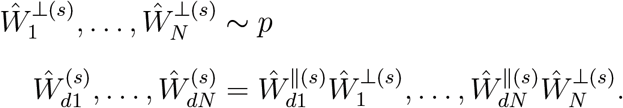

This modification turns ALDEx2 into a specialized type of model called a Scale Simulation Random Variable (SSRV) (Nixon et al., 2023).

### Simple Scale Models for DA/DE Analysis

While scale models are flexible and can be arbitrarily complex, they do not need to be. Especially for DA/DE analysis, the structure of the LFC estimand (*θ*_*d*_) can simplify model specification. Nixon et al. (2023) proved that for LFC estimation, scale models only need to be specified up to a global constant *c* defined by 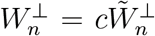. This result implies that researchers designing scale models for DA/DE analysis only need to be concerned with how the scale might change between systems. Moreover, for LFC estimation, those authors showed that it often suffices to specify a scale model for a single real-valued quantity *θ*^*⊥*^ called the *Log-Fold-Change in Scales*: defined by

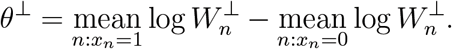

That is, in many cases, it is sufficient only to model how the average scale might change between conditions. The scale models used in this manuscript use one or both simplifications.

### A Default Scale Model for ALDEx2

To ease adoption of the updated ALDEx2 software suite, we developed a scale model that considered potential error in the default CLR normalization. Yet, we avoid using Equation (1) for this task as, due to the law of large numbers, that model asymptotically assumes that the LFC of scales is equal to the CLR estimate with zero uncertainty (zero variance). Instead, we defined an alternative model with better asymptotic performance that more naturally mimics the linear modeling capabilities of ALDEx2. We present the model in its full form in Supplementary Section S.3. For DA/DE analyses, the scale model simplifies to

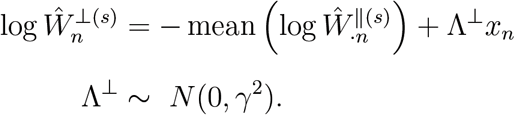

As in Equation (1), this scale model reduces to the CLR normalization when *γ* = 0 and models error in that assumption for any value of *γ >* 0.

The parameter Λ^*⊥*^ represents systematic error in the CLR estimated difference in scales between conditions. More concretely, using Equation (3) we calculate that the CLR normalization corresponds to an assumption that

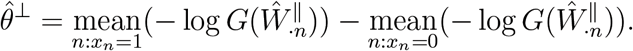

The parameter Λ represents potential error in this relationship; the true log-fold-change in scales (*θ*^*⊥*^) is given by 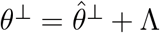. Considering the distribution of Λ, this implies a model 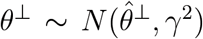. When choosing *γ*, one should consider 95% probability intervals of this normal model. According to this model, there is a 95% probability that the true difference in scales between conditions is within a factor of 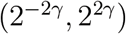 of the CLR estimate 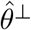. Based on this interpretation, we recommend *γ* = 0.5 as a reasonable choice: at this value, we consider up to 2-fold errors in the CLR estimate of *θ*^*⊥*^ with 95% probability. Alternatively, rather than interpreting Λ in terms of errors in the CLR estimate, it can be interpreted directly in terms of the log-fold-change of scales. According to the model, there is a 95% probability that the true difference in scales between conditions is within the range 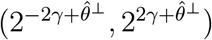. See Supplemental Section S.3 and S.4 for further details on interpretation of the default scale model and advice for choosing *γ*.

### Updates to the Summarization of p-values in ALDEx2

Upon introducing scale models in ALDEx2, we identified a slight error in how ALDEx2 had previously summarized p-values over the *S* posterior samples. While unlikely to cause issues in the normalization-based ALDEx2, this error grew problematic when we introduced scale uncertainty. In Supplementary Section S.2, we illustrate this problem and describe a solution of summarizing p-values from two one-sided hypothesis tests rather than from a single two-sided test.

## Data Analysis Details

For all data analyses, p-values reported for ALDEx2 refer to Benjamini-Hochberg corrected p-values of the null hypothesis *θ*_*d*_ = 0 using the Welch’s t-test and based on 1,000 Monte Carlo replicates. The exception is our analysis of the Gierliński et al. (2015) RNA-seq study presented in Supplementary Section S.6, where we only used 500 Monte Carlo replicates to accommodate the larger data size. For all analyses, DESeq2, edgeR, limma-voom, and baySeq were fit using recommended defaults. For edgeR, we report the results of the exact test. For clarity, logarithms in the following sections were computed in base 2 to be consistent with ALDEx2.

### Mock experiment simulation details

The true abundance of 20 microbial taxa were simulated from 2*N* communities equally split between pre-antibiotic (*x*_*n*_ = 0) and postantibiotic (*x*_*n*_ = 0) conditions. Simulations used the following Poisson model:

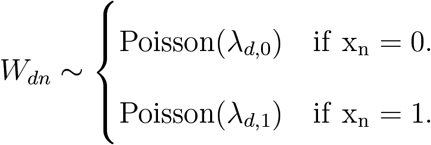

To simulate a narrow-spectrum antibiotic, 16 of the 20 taxa were specified with *λ*_*d*,0_ = *λ*_*d*,1_ (not differentially abundant). Of these 16 non-differentially abundant taxa, 4 had *λ*_*d*_ = 4, 000, 3 had *λ*_*d*_ = 500, and 9 had *λ*_*d*_ = 400. The four taxa that were differential abundant were *d* = *{*3, 4, 15, 20*}*. For those taxa, *λ* was set as: *λ*_(3,4,15, and 20);0_ = *{*4, 000, 4, 000, 400, 400*}* and *λ*_(3,4,15, and 20);1_ = *{*3, 000, 2, 000, 200, 100*}*. Based on these values, the CLR estimate of the LFC of scales was 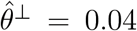 whereas the true value was *θ*^*⊥*^ = *−*0.18. That is, the CLR assumption implies a slight increase in scales after antibiotic administration; the truth is a moderate decrease in scales after antibiotic administration. Sequencing-based loss of scale information was simulated via multinomial resampling:

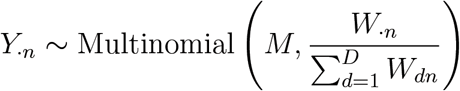

with a sequencing depth *M* = 5, 000.

The Informed model was constructed under the assumption that antibiotic administration resulted in a 10% decrease in the total microbial load between conditions:

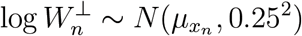

where 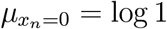 (pre-antibiotic) and 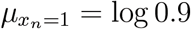 (post-antibiotic).

### SELEX Reanalysis

The SELEX experiment is detailed in McMurrough et al. (2014). Preprocessed data from this experiment was obtained from the ALDEx2 Bioconductor package. This data contains 1,600 possible sequence variants measured in 14 samples equally split between the selected and non-selected conditions. True positives were identified based on subsequent validation experiments detailed in McMurrough et al. (2014).

For this study, the CLR estimate for the log-fold-change of scales 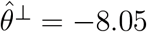 corresponds to an assumption that the average scale in the selected condition is 265 times higher than in the non-selected condition. The default scale model at *γ* = 0.5 expresses 95% certainty that the average scale in the selected condition is between 130 to 530 times larger than in the non-selected condition. At *γ* = 5, the default scale model expresses 95% certainty that the average scale in the selected condition is between 0.25 to 272,000 times larger than in the non-selected condition. In reality, our true beliefs lie in between these two values of *γ*. We selected *γ* = 0.5 as a reasonable default value and *γ* = 5 to highlight performance under unreasonably large amounts of uncertainty.

Data resampling was performed in triplicate for each sample size to address randomness in the resampling process.

### Vandeputte Reanalysis

Data was obtained from the European Nucleotide Archive with accession code PRJEB21504. A sequence variant table using the DADA2 software processed the raw data, following the software vignette and recommended defaults (Callahan et al., 2016). Fastq files were filtered with the filterAndTrim function setting maxEE to 4 as described in online vignettes. We used all default parameters to learn the error rates, run the core DADA2 algorithm, and merge sequence pairs. The consensus method removed chimeras. We used the RDP classifier to assign taxonomy (Wang et al., 2007), then retained genera present in at least 20% of samples for analysis. QMP was applied by using R code available at https://github.com/raeslab/QMP. The resulting matrix was treated as an estimate of *W* . A small pseudo-count was added (0.5) prior to log-transformation to mitigate numerical issues associated with taking the logarithm of zero. Two-sided Welch’s t-tests were applied to assess significance between CD patients and health controls for each genera. Resulting p-values were adjusted using the Benjamini-Hochberg procedure.

If we assume the flow cytometry measurements are error free, then the microbial load in CD patients decreases by 65% compared to healthy controls (*θ*^*⊥*^ = *−*1.49). However, extended results presented by Vandeputte et al. (2017) also suggest that these measurements can have substantial variability leading us to the following (Gold Standard) scale model:

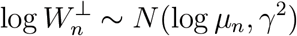

where *μ*_*n*_ denotes the measured flow-cytometry cell count for sample *n*, and *γ*^2^ is the variance of the measurement noise. We chose *γ*^2^ based on based on Extended Data File 5 of Vandeputte et al. (2017) which summaries the mean and standard deviation cell counts from technical replicates of 40 different biological samples. A Taylor expansion was used to estimate standard deviations on log-scale based on the reported means and standard deviations of cell counts. Conservatively, we chose a value of *γ*^2^ = 0.70 which corresponds to the maximum estimated log-scale variance from the 40 samples studied. This model expresses 95% certainty that the average scale in Crohn’s disease patients is between 13% and 94% of the average scale in the healthy controls. Figure 4 depicts the sensitivity of results to this choice of *γ*^2^.

An Informed scale model was designed based on visual inspection of Figure 2 of Sarrabayrouse et al. (2021). Based on that figure, we estimated an average of 1.5 *×* 10^12^ cells per gram of feces for healthy controls compared to 1.0 *×* 10^12^ cells per gram of feces for CD patients. Combined with estimates of uncertainty obtained from that figure, we designed a scale model which reflects an assumption of an approximately 30% decrease in microbial load in CD compared to health:

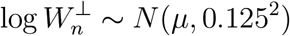

where *μ* = log(1) if the sample was from the control condition and *μ* = log(0.7) if the sample was from the CD condition (*θ*^*⊥*^ = *−*0.52). This model expresses 95% certainty that the average scale in the Crohn’s disease patients is between 17% and 41% lower than the average scale in the healthy patients.

## Data and Software Availability

All changes discussed in this manuscript are available as of package version 1.34.0, which is in the version 3.18 Bioconductor release or later. With the release, a vignette “Incorporating Scale Uncertainty into ALDEx2” reviews the new capabilities of ALDEx2 and describes how to use the default scale model, or any user-specified scale model. This vignette and accompanying documentation also describe other features that have been added to facilitate scale model-based analyses.

All data and code needed to reproduce the analyses in this article are available at https://github.com/michellepistner/scale-in-aldex2.

## Acknowledgments and Funding Sources

We thank Rachel Silverman and Steve Nixon for their manuscript comments and Yen Duong for her professional editing services. JDS and MPN were supported in part by NIH 1R01GM148972-01. GBG declares no specific funding for this work.

## Supporting Information

### Supplement

Contains derivations for the TSS normalization, information on the updates to the testing procedure, a description of ALDEx2 as a linear model, further details on selecting *γ*, details on sensitivity analyses, an expanded analysis of an RNA-seq data set, and Supplemental Figures S.1-S.2.

## Footnotes

1 We express this as a model for log 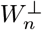 rather than 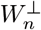 so that it can be expressed as a normal distribution rather than the lesser known log-normal while still restricting 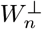 only to take on positive values.

