## Supplemental materials for "Beyond Normalization: Incorporating Scale Uncertainty in Microbiome and Gene Expression Analysis"

### S.1 Implied Scale Assumption of the TSS Normalization

In this section, we show that Total Sum Scaling normalization (TSS) leads to the implicit assumption that scale remains constant between conditions. As in *Methods*, let $W_{dn}$ denote the amount of entity $d$ in sample $n$. Recall that this amount can be decomposed as the product of the proportional abundance of entity $d$, $W^{\parallel}_{dn}$, and the total scale of the system $n$, $W^{\perp}_{n}$:

$$W_{dn} = W^{\parallel}_{dn} W^{\perp}_{n}.$$

The log-fold-change (LFC) of entity $d$ can be decomposed into two additive components, one depending only on $W^{\parallel}$ and one on $W^{\perp}$:

$$\begin{aligned}
\theta_d &= \underset{n:x_n=1}{\text{mean}} \log W^{\parallel}_{dn} W^{\perp}_{n} - \underset{n:x_n=0}{\text{mean}} \log W^{\parallel}_{dn} W^{\perp}_{n} \\
&= \underbrace{\underset{n:x_n=1}{\text{mean}} \log W^{\parallel}_{dn} - \underset{n:x_n=0}{\text{mean}} \log W^{\parallel}_{dn}}_{\theta^{\parallel}_{d}} + \underbrace{\underset{n:x_n=1}{\text{mean}} \log W^{\perp}_{n} - \underset{n:x_n=0}{\text{mean}} \log W^{\perp}_{n}}_{\theta^{\perp}}
\end{aligned}$$

In TSS, the observed data $Y$ is normalized and used as estimates of $W^{\parallel}$, yet LFCs are calculated directly from these normalized estimates:

$$\begin{aligned}
\hat{W}^{\parallel}_{dn} &= \frac{Y_{dn}}{\sum_{d=1}^{D} Y_{dn}} \\
\hat{\theta}_d &= \underset{n:x_n=1}{\text{mean}} \hat{W}^{\parallel}_{dn} - \underset{n:x_n=0}{\text{mean}} \hat{W}^{\parallel}_{dn}.
\end{aligned}$$

From $\hat{\theta}_d$ above, it is clear that this is simply an estimate of the compositional term $\theta^{\parallel}$. In other words, using TSS normalized data for LFC estimation is equivalent to an implicit assumption that $\theta_d = \theta^{\parallel}_d$ or equivalently that $\theta^{\perp} = 0$. The assumption that $\theta^{\perp} = 0$ occurs when the average scale in one condition is exactly equal to the average scale in the other, i.e.:

$$\underset{n:x_n=1}{\text{mean}} \log W^{\perp}_{n} = \underset{n:x_n=0}{\text{mean}} \log W^{\perp}_{n}.$$

### S.2 An updated hypothesis test for ALDEx2

While adding scale to ALDEx2, we discovered an error in how ALDEx2 had been summarizing p-values over the $S$ simulation samples. While this error could lead to counter-intuitive results, we expect that it would rarely, if ever, cause issues in a normalization-based ALDEx2, yet the error may become problematic when scale models are added.

For each entity $d$, ALDEx2 computes an average p-value across the $S$ Monte Carlo replicates. Originally, a p-value, $p_{ds}$, was computed for each entity $d \in 1, ..., D$ and Monte

Carlo sample $s \in \{1, ..., S\}$ using a two-sided alternative, i.e. in the case of a two sample t-test:

$$p_{ds} = \Pr(|T| > |t_{ds}| \ \mid H_0) \tag{1}$$

$$p_d = \frac{1}{S}\sum_{s=1}^{S} p_{ds} \tag{2}$$

where $t_{ds}$ denotes the observed value of the test statistic and $T$ denotes the the distribution of the test statistic. Since ALDEx2 is averaging over multiple p-values, this specification can be problematic if the signs of the test statistic vary over the $S$ replicates for a given entity $d$. To see this concretely, consider a simple example where there are two conditions with 10 samples per condition ($N = 20$). Suppose that, for a given entity, two Monte Carlo samples are drawn, resulting in the test statistics $t_{d\cdot} = \{-3, 3\}$. The two-sided p-values are $p_{d\cdot} = \{0.008, 0.008\}$ (18 degrees of freedom) which corresponds to $p_d = 0.008$. At face value, the aggregated p-value would suggest that there is a significant effect, yet the two Monte Carlo samples disagree on the sign of the effect.

This problem is well-known in the statistical literature of combining p-values from multiple hypothesis tests (Oosterhoff, 1969). It can largely be avoided by careful selection of the tested hypotheses. In line with recommendations in the literature (Oosterhoff, 1969), we propose a slight modification of the original approach in ALDEx2 and average over p-values obtained from one-sided tests ($p_{ds}^{\text{mod}}$) instead of two-sided tests:

$$p_{ds}^{\text{mod}} = Pr(T > t_{ds} \ |H_0) \tag{3}$$

$$p_d^{\text{mod}} = \frac{1}{S}\sum_{s=1}^{S} p_{ds}^{*} \tag{4}$$

$$p_d^{\text{mod}} = 2 \times \min(p_d^{*}, 1 - p_d^{*}). \tag{5}$$

where $T$ is defined as before and $p_d^{\text{mod}}$ denotes the p-value obtained from this modified procedure for entity $d$. In words, we compute the one-sided p-value for a right-tailed alternative for all Monte Carlo samples ($p_{ds}^{\text{mod}}$), average over all Monte Carlo samples ($p_d^{\text{mod}}$), and select the direction that leads to the lowest overall p-value ($p_d^{\text{mod}}$) after multiplying by a factor of two to convert two a two-sided p-value. Equation (5) holds due to the fact that the left-tailed and right-tailed p-value must sum to one due to complimentary events.

Practically, the new procedure requires that each Monte Carlo replicate offers evidence towards both the magnitude and direction for an entity to be returned as significant. For example, $p_d^{\text{mod}} = 1$ in our toy example. More generally, the modified p-value ($p_d^{\text{mod}}$) will be larger or equal to the original p-value ($p^d$) produced by ALDEx2. If all Monte Carlo replicates agree on sign, the two will be equal ($p_d^{\text{mod}} = p^d$), otherwise $p_d^{\text{mod}} > p^d$.

### S.3 ALDEx2 as a Linear Model

While the results in the main text were described for differential abundance or expression, they apply more generally to linear models. Note that differential abundance and expression is equivalent to a linear model with an intercept and a binary covariate denoting the condition.

To view the results of the main text as a linear model, let $\hat{\theta}_{dr}$ denote the estimated linear relationship between covariate $r$ and entity (taxa or gene) $d$: i.e., an estimate of the parameter in a linear model of the form $\log W_{dn} = \alpha_d + \sum_r \theta_{dr} X_{rn} + \epsilon_{dn}$. In words, $\theta_{dr}$ captures the association between changes in the $r$-th covariate and changes in the amount of entity $d$. When viewed as a linear model, estimates produced by ALDEx2 suffer from the same phenomenon described in the main text: $\hat{\theta}_{dr}$ captures the association between changes in the covariate $r$ and changes in the *normalized* amount of entity $d$. This leads to the same phenomenon of unacknowledged bias described in the main text.

Procedurally, ALDEx2 remains virtually identical to the method for differential abundance and expression with one exception: for each taxa $d$ and sample $s$, the t-test is replaced by a linear model with mean (i.e., expectation, $\mathbb{E}$):

$$\mathbb{E}[\log \hat{W}_{dn}^{(s)}] = \theta_{d0} + \theta_{d1}X_{1n} + \cdots + \theta_{dR}X_{Rn},$$

resulting in estimates $\hat{\theta}_{dr}^{(s)}$ and p-values $p_{dr}^{(s)}$ corresponding to the null hypothesis $\theta_{dr} = 0$ (using a t-test or Wald test). Corrected p-values using either the Holm or Benjamini-Hochberg procedure are also returned. These estimates and p-values are summarized over the $S$ samples in an analogous manner to the procedure described for differential abundance or expression analysis.

#### S.3.1 Default Scale Model in Linear Models

In the case of linear models, the new default scale model is:

$$\log \hat{W}_{n}^{\perp(s)} = -\log \phi\left(\hat{W}_{\cdot n}^{\parallel(s)}\right) + \sum_{r=1}^{R} \Lambda_r^{\perp} x_{rn}$$

$$\Lambda_r^{\perp} \sim N(0, \gamma^2).$$

where $x_{rn}$ denotes the user specified covariates $X_{rn}$ re-scaled so that continuous covariates have mean zero and standard deviation 1, and binary covariates are coded as 0 or 1. This scaling ensures the resultant model is invariant to arbitrary recodings of the covariates, e.g., it is invariant to whichever binary encoding is used for $X$.

Since the default scale model is an extension of the one presented in the main text for differential abundance and expression, it retains the design features described previously: namely, it requires only a single input and will reduce type-I error rates compared to the CLR normalization. In addition, this scale model has a third property for linear models that we find appealing: the term $\sum_{r=1}^{R} \Lambda_r^{\perp} x_{rn}$ scales with the number of covariates $R$. The rational is as follows: if each covariate could change the scale of the system under study (e.g., a drug could kill microbes), then our uncertainty in the system scale should also scale with the number of covariates.

### S.4 Advice on Choosing $\gamma$

For simplicity, we state this advice in terms of the default scale model for differential abundance. Extending this intuition to the default scale model for linear models is straightforward. As discussed in *Methods*, the parameter $\gamma$ controls the amount of uncertainty in the scale assumption implied by the CLR normalization. Similarly, $\Lambda^{\perp}$ is a factor representing how the scale changes between biological conditions with larger values of $\gamma$ corresponding to more scale uncertainty. Thus, the default scale model assumes that, with 95% certainty, the log-fold-change of scales between conditions is between $(2^{-2\gamma+\hat{\theta^{\perp}}}, 2^{2\gamma+\hat{\theta^{\perp}}})$.

This interpretation can be used to find a reasonable value of $\gamma$ in a given context. For example, consider the simulation presented in the main text. Suppose we know the antibiotic is narrow spectrum and expect that scale changes minimally between conditions (most taxa are not affected): we expect that the antibiotic is unlikely to kill more than 20% of microbial load. We can use the Dirichlet samples to get an approximation of the CLR estimate ($\hat{\theta}^{\perp} = 0.04$), which means the CLR assumption is estimating a slight increase in microbial load after antibiotic treatment. As this seems counter-intuitive, if we want to still use the default scale model we should choose a suitably large value of $\gamma$ to cover our beliefs and account for this bias in $\hat{\theta}^{\perp}$. Equating our assumption that the antibiotic is unlikely to kill more than 20% of the microbial load with the lower bound of our 95% probability region of the default scale model, we can calculate the value of $\gamma$ by solving the equation:

$$2^{-2\gamma+0.04} = 0.8, \tag{6}$$

yielding a value of $\gamma = 0.18$:

$$\gamma = \frac{\log_2 0.8 - 0.04}{-2} \tag{7}$$

$$\gamma = 0.18. \tag{8}$$

### S.5 Sensitivity Analyses

To facilitate the choice of $\gamma$, we developed and implemented a special sensitivity analysis for the default scale model. Simply, this sensitivity analysis repeats our method over a user-specified range of $\gamma$, allowing for an easy comparison of results over different scale assumptions. This algorithm was made efficient using the specialized memorization algorithm developed in Nixon et al. (2023), and the sensitivity analysis is implemented in the `aldex.senAnalysis` function in the `ALDEx2` package. A special plotting function is available to visualize these results (`plotGamma`); an example of the visualized output is in Figure S.1.

To show how these visualizations can be helpful, consider Figure S.1. There is a large cluster of taxa that are only significant (black) for tiny values of $\gamma$. This cluster contains all of the false positives. Including even small amounts of scale uncertainty (choosing small but non-zero $\gamma$) is enough to remove these false positives. Even if we did not know which were true versus false positives, seeing a cluster of taxa so sensitive to even slight amounts of error in scale assumptions should be sufficient to allow researchers to avoid drawing conclusions about these taxa and would suggest that $\gamma$ should be set to be greater than 0.1.

In addition, sensitivity analyses can eliminate the need to choose $\gamma$ and facilitate a novel, reproducible, and transparent reporting form. Rather than specifying a value of $\gamma$ a priori, a researcher would report the maximum value of $\gamma$ under which a result holds, e.g., based on Figure S.1, taxa 20 is differentially decreased after antibiotic treatment ($p < 0.05$) so long as $\gamma < 0.70$. Overall, we would expect that such a reporting would enhance the transparency and reproducibility of sequence count data analyses, allowing readers to interpret the robustness of reported conclusions themselves.

### S.6 Analysis of an RNA-Seq Dataset

We reanalyzed an RNA-seq study originally published in Gierliński et al. (2015) and reanalyzed in Schurch et al. (2016) which contained 48 biological replicates from either wild-type (WT) or a SNF2-knockout (SNF2) strain of *Saccharomyces cerevisiae*. Raw reads for this data set was obtained from the European Nucleotide Archive repository (ENA, accession code PRJEB5348, `http://www.ebi.ac.uk/ena/data/view/ERX425102`) and processed as in Schurch et al. (2016). Aggregated data is also available at `https://github.com/ggloor/datasets`. To mimic Schurch et al. (2016), the data set was restricted to only those replicates that passed cleaning checks described in Gierliński et al. (2015), resulting in 44 $\Delta$snf2 biological replicates and 42 wild-type replicates. Entities were filtered such that all entities with zero counts across all of the restricted replicates were removed, resulting in 6,327 entities. Full experimental details can be found in Gierliński et al. (2015) and Schurch et al. (2016).

Prior analyses of these data have shown that standard differential expression tools typically report about 70% of genes as being differentially expressed at the full data size (Schurch et al., 2016). These results seem implausible. While SNF2 is a key gene involved in chromatin remodeling (Peterson and Tamkun, 1995), we think it unlikely that this single gene knockout leads to change in the expression of over 70% of expressed genes. Instead, we suspect that this might be a problem of unacknowledged bias, especially as prior authors have noted that the positive identification rate drastically rises as a function of sample size (Schurch et al., 2016). While we do not know which genes are truly differentially expressed within this study, we can investigate our hypothesis by quantifying the sensitivity of differential expression to modeling assumptions about scale. Central to our argument is the idea that a scientific conclusion is suspect if it only holds under a very restrictive set of unrealistic modeling assumptions.

We compared the results from ALDEx2 using two different scale models: the default scale model and an *informed* scale model built based on consideration of the experimental design. This model was specified using the results of Yoshikawa et al. (2011) which suggested a 10% decrease in the SNF2 deficient strains compared to wild-type. Echoing this belief, we used this as a basis for our informed scale model:

$$\log(W_{\text{n}}^{\perp(s)}) \sim N(\mu, \gamma^2) \tag{9}$$

where $\log(W_{\text{n}}^{\perp(s)})$ indicates the $s^{th}$ draw from the scale model for sample $n$, $\mu = \log(1)$ for the wild-type strain, and $\mu = \log(0.9)$ for the SNF2 strain. Within the context of this study, the CLR assumption corresponds to an implicit assumption that total RNA expression in the WT strain is about 20% higher than in the SNF2 strain. Note that the assumed direction is the same as the custom scale model. For both the default scale model and the user-specified model, we set $\gamma = \{0, .25, .5, .75, 1, 1.25, 1.5, 1.75, 2\}$, sampled 500 Monte Carlo replicates, applied Welch's t-test, and selected significant entities according to a corrected p-value threshold of 0.05. The results are summarized in Figure S.2.

Echoing the results of Schurch et al. (2016), at the full data size, ALDEx2 with the CLR assumption (default scale model with $\gamma = 0$) finds that approximately 68% of genes are differentially expressed between conditions. As hypothesized, this result drops dramatically with even low levels of scale uncertainty. For example, at $\gamma = 0.25$, this figure drops from 68% to just 12%. Similar results are seen for the ALDEx2 with the informed scale model. For the smallest value of $\gamma$, the two scale models generally agree, but, for larger values of $\gamma$, the informed scale model returns more genes as significant than the default scale model, representing potential loss of power if the default scale model is biased. Additionally, we note that the knockout gene (YOR290C) is significant for either scale model for all values tested and has the largest effect size for both models when $\gamma = 2$.

Overall, these sensitivity analyses suggest that less than 70% of genes are differentially expressed when even small amounts of scale are included in the model, and that prior reports may have been biased due to error caused by inappropriate assumptions about scale.

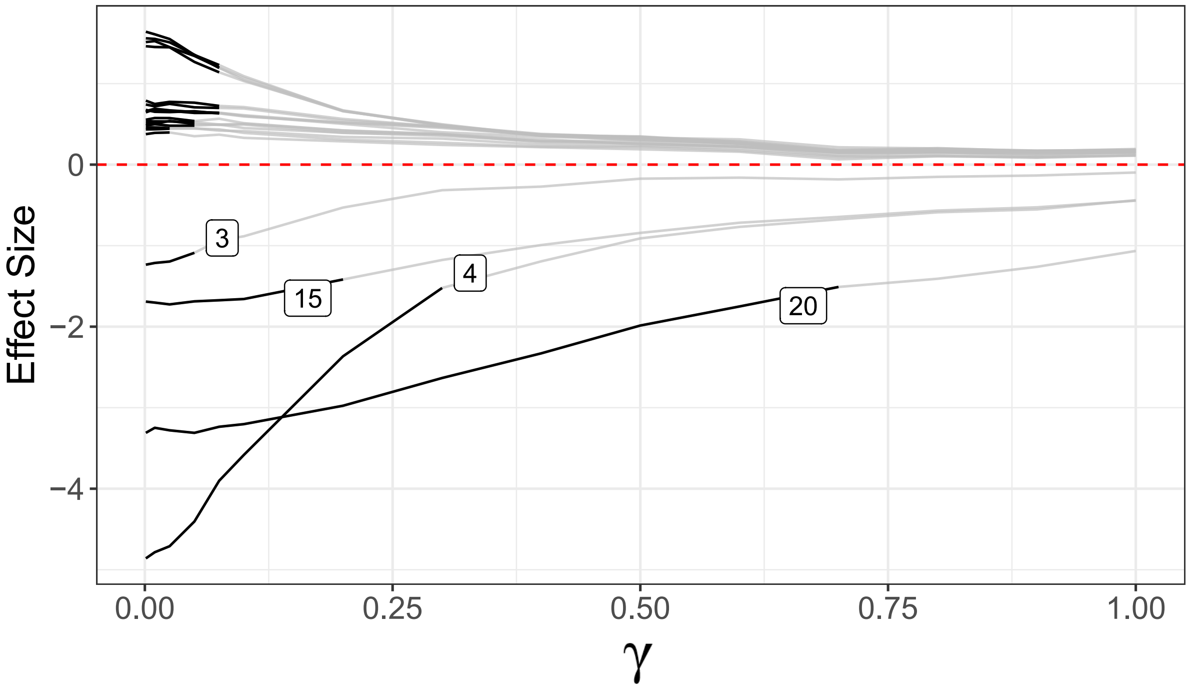


Figure S.1: **Sensitivity analysis for ALDEx2 with varying levels of $\gamma$.** We conducted a sensitivity analysis of the simulation data using the default scale model and fitting procedures as described in *Methods*, varying the amount of uncertainty added to the default scale model ($\gamma \in [0, 1]$). At each value of $\gamma$ and for each entity, we computed the log fold change (LFC) for each Monte Carlo sample and calculated the mean and standard deviation across these samples. Then, we estimated a standardized value of the LFC by dividing the mean by its corresponding standard deviation. Standardized LFCs as a function of $\gamma$ were plotted, and lines were marked in black if the p-value for that entity was less that 0.05. Even as $\gamma$ increases, several of the true positive results remain, e.g. taxa 4, 15, 20.

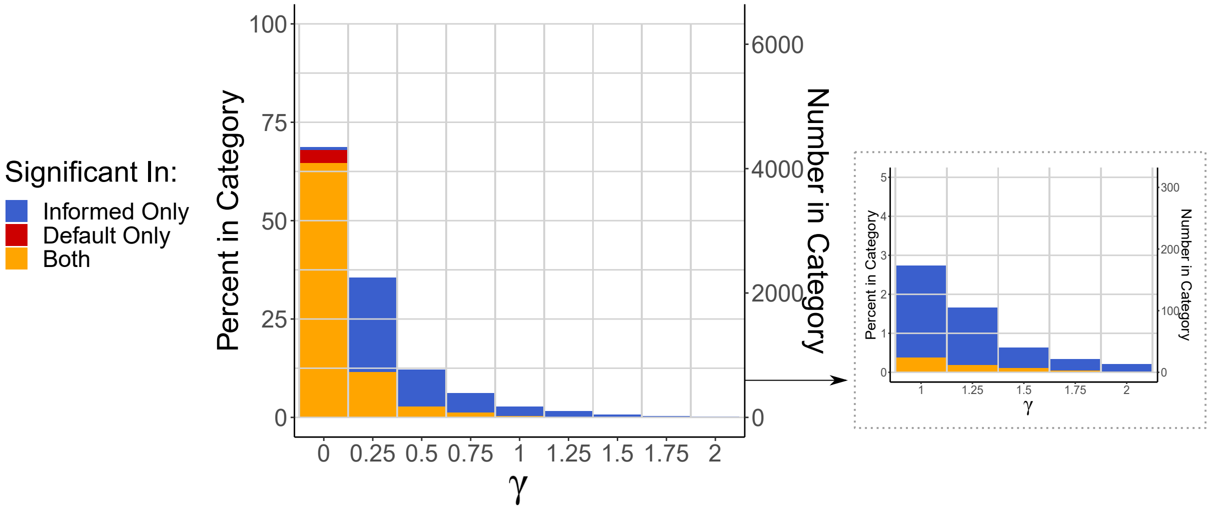


Figure S.2: **A sensitivity analysis for the RNA-seq data.** We analyzed the RNA-seq data set using the updated ALDEx2 model as outlined in Section S.6. We tested two scale models, the default scale model and the Informed model, varying the amount of uncertainty added ($\gamma$). For each model and value of $\gamma$ tested, we recorded which entities were significant and compared these results to the opposing scale model. As scale uncertainty increases, the number of positives drops regardless of scale model (e.g., approximately 65% of entities are positive at $\gamma = 0$ compared to 1.5% of entities at $\gamma = 0.5$ for the default scale model).
